# Dynamic multimodal survival prediction in multiple myeloma integrating gene expression, longitudinal laboratories, and treatment history

**DOI:** 10.64898/2026.03.30.715136

**Authors:** Shangru Jia, Artem Lysenko, Keith A Boroevich, Alok Sharma, Tatsuhiko Tsunoda

## Abstract

Prognostic stratification in multiple myeloma (MM) relies on staging systems that assign patients to fixed categories at diagnosis and discard the temporal information that accumulates during treatment. We developed a dynamic multimodal framework that predicts residual overall survival using observation windows ranging from 1 to 18 months post-diagnosis. The model integrates DeepInsight-transformed gene expression representation, longitudinal laboratory measurement trajectories across 10 analytes, and treatment history for three drug classes through an adaptive fusion mechanism that accounts for missing clinical observations. On the MMRF CoMMpass cohort (*n* = 752), five-fold cross-validation yielded a concordance index (C-index) of 0.773 ± 0.024 and a time-dependent AUC at a 1-year prediction horizon (tdAUC_1yr_) of 0.789 ± 0.021, outperforming all evaluated baseline methods including DeepSurv (0.633 ± 0.095) and random survival forests (0.636 ± 0.024) on matched cross-validation splits. Modality ablation identified longitudinal laboratory measurements as the strongest individual contributor (C-index 0.693); the DeepInsight spatial encoding of gene expression yielded higher discrimination than a multilayer perceptron (MLP) baseline operating on the same features (0.624 vs. 0.596). Kaplan–Meier analysis showed significant prognostic group separation at all primary landmarks (log-rank *p <* 0.001; hazard ratios 3.46–3.93). A distilled student model retaining only the DeepInsight representation and five baseline clinical features achieved C-index 0.672 and tdAUC_1yr_ 0.740 on an independent microarray cohort (GSE24080, *n* = 507) without retraining. Interpretability analysis identified prognostic associations consistent with established myeloma biology, including ubiquitin-proteasome pathway genes, endoplasmic reticulum stress markers, and Interferon Alpha Response pathway enrichment.

## 1. Introduction

Multiple myeloma (MM) is a clonally heterogeneous plasma cell malignancy accounting for approximately 10% of hematologic cancers, with overall survival spanning months to over a decade among newly diagnosed patients [1, 2]. The International Staging System (ISS) and its revisions (R-ISS, R2-ISS) combine serum *β*_2_-microglobulin (*β*_2_M), albumin, lactate dehydrogenase (LDH), and high-risk cytogenetic aberrations into ordinal stages that remain the clinical standard for risk classification [3, 4, 5, 6]. However, these systems share a fundamental constraint: they assign patients to fixed categories at diagnosis and do not incorporate the longitudinal biomarker dynamics or treatment context that accumulate over the course of therapy.

Computational approaches to survival prediction in MM have pursued several directions. Random forest models combining clinical and RNA-seq features on the CoMMpass cohort achieved C-indices near 0.78 [7], and the transformer-based SCOPE framework addressed joint event prediction across disease phases [8]. Deep learning survival methods have evolved from single-time-point architectures such as DeepSurv [9] and DeepHit [10] to longitudinal formulations including Dynamic-DeepHit [11]. In parallel, multimodal fusion methods in computational pathology— including MCAT [12], Pathomic Fusion [13], PORPOISE [14], and pathway-aware multimodal transformers such as SurvPath [15]— have demonstrated that cross-modal integration can improve survival discrimination [16]. However, these architectures operate at fixed time points on histopathology–transcriptomics pairs and do not address the temporal, multi-source data structure characteristic of longitudinal hematologic care.

A distinguishing feature of MM management is its prolonged, multi-phase treatment course, spanning induction, transplant eligibility assessment, consolidation, and maintenance over months to years [1, 2]. During this period, prognosis-relevant information accumulates continuously. Biomarker trajectories—including free light chain kinetics, serum M-protein response, shifts in *β*_2_M and LDH—reflect treatment response, residual disease burden, and emerging clonal dynamics in ways that a single diagnostic snapshot cannot capture. In practice, clinicians reassess prognosis at defined milestones such as induction completion, transplant decision, and maintenance initiation. Yet most existing computational models in MM are trained on features captured at a single time point and therefore cannot be easily updated as new observations become available. This motivates development of a prediction framework that incorporates time-varying clinical data and revises prognosis estimates dynamically over the treatment course.

Two methodological ideas motivate the present study. First, the landmark prediction framework [17] enables dynamic risk estimation at successive time points using all previously observed data; we extend this paradigm by processing full longitudinal sequences through temporal encoders rather than treating covariates as fixed summary values at each landmark. Second, gene expression profiling has established prognostic value in MM through signatures such as GEP70 [18] and EMC-92 [19], but incorporating high-dimensional expression data into predictive models remains challenging because of the curse of dimensionality. DeepInsight [20] addresses this problem by restructuring tabular features into two-dimensional image representation that preserve local feature relationships, enabling convolutional neural networks (CNNs) to exploit spatial co-expression structure. Although this transformation has been applied to drug response prediction [21], it has not been explored in survival modelling. Separately, knowledge distillation [22] has been used to compress survival models [23] but has not been combined with multimodal dynamic prediction in hematologic malignancies. In MM, external cohorts frequently provide only baseline expression data without longitudinal laboratory or treatment records. Compressing a multimodal model into a reduced-input student could facilitate deployment in external settings where not all data modalities are available.

Figure 1 summarizes the overall framework. The model integrates three data modalities: gene expression image representation converted by DeepInsight, longitudinal laboratory trajectories, and treatment history—through gated fusion conditioned on observation reliability indicators, producing risk estimates from any observation window within the first 18 months post-diagnosis. We evaluate the framework on the MMRF CoMMpass dataset using five-fold cross-validation, benchmark it against standard survival methods, validate it externally on the GSE24080 cohort using a distilled student model, and provide interpretability analyses linking model predictions to myeloma biology.

**Figure 1.**
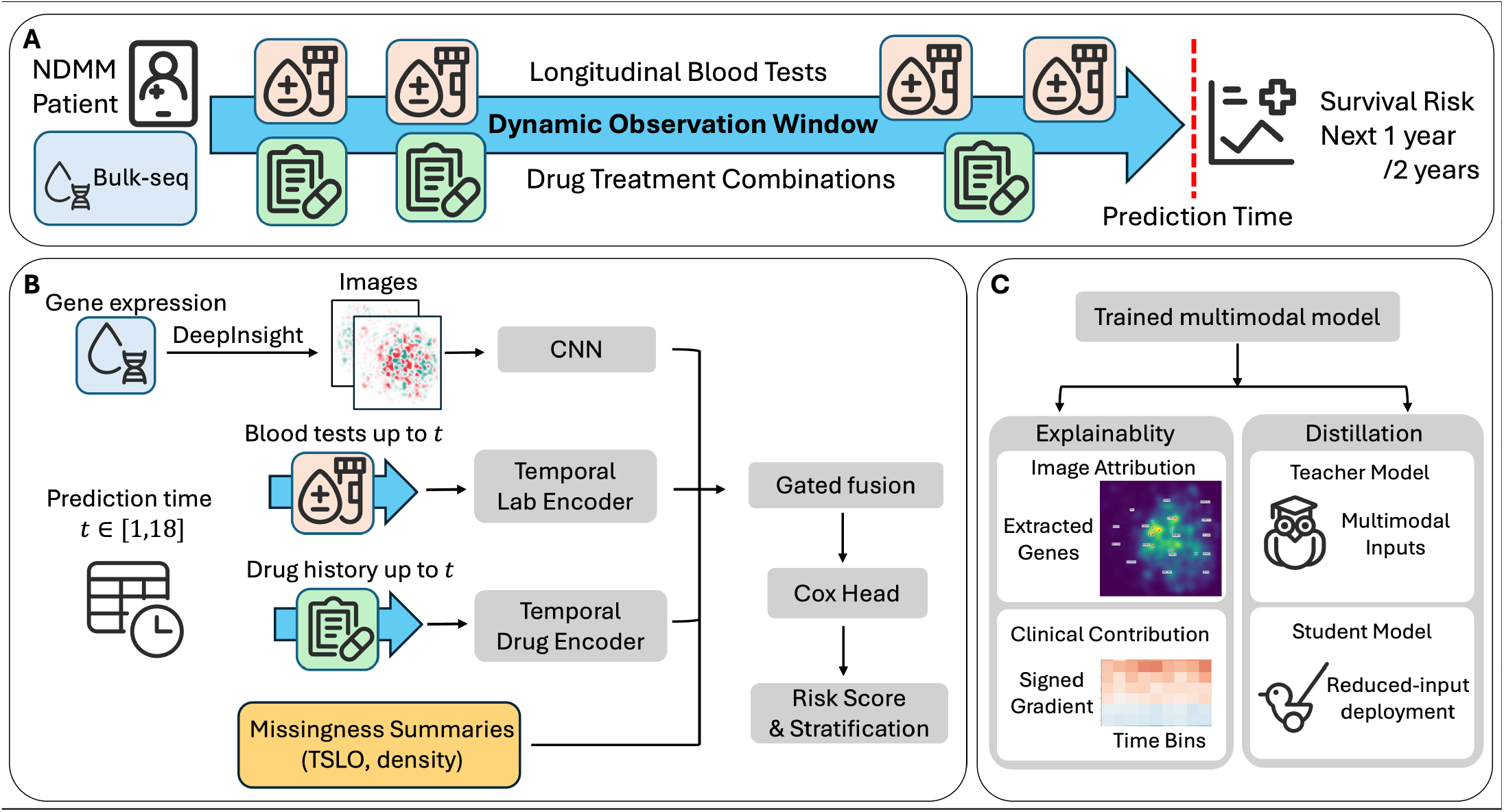
Study overview. (A) A newly diagnosed MM patient accumulates longitudinal blood tests and drug treatment records over a dynamic observation window; at any prediction time t ∈ [1, 18] months, the model estimates residual survival risk using all data observed up to t, combined with a baseline DeepInsight gene expression image. (B) Model architecture: modality-specific encoders feed into missingness-aware gated fusion; a Cox PH head outputs the log-hazard risk score, with auxiliary per-modality heads providing gradient supervision. (C) Downstream applications: interpretability and teacher–student distillation for reduced-input external deployment.

## 2. Materials and Methods

### 2.1 Study design and cohort construction

The primary development cohort was drawn from the MMRF CoMMpass study (NCT01454297), a prospective longitudinal study of newly diagnosed MM patients with paired baseline bulk RNA-seq, serial laboratory measurements, and treatment records [24, 25]. Patients were included if they had (i) available baseline RNA-seq expression data, (ii) at least one post-diagnosis laboratory observation, and (iii) minimum follow-up data sufficient to contribute to at least the earliest landmark (*t* = 1 month). After applying these criteria, 752 patients were retained. Of these, 752 patients were split by stratified random sampling on event status into a development set of 624 (124 deaths, 19.9% event rate; overall survival range 41– 1984 days) and a held-out validation set of 128 (24 deaths, 18.8%), prior to any preprocessing. The development set was used for all model training, hyperparameter tuning, and cross-validation. The held-out set was withheld from all stages of model development; no hyperparameter selection, checkpoint selection, or architecture decisions were informed by this subset. The selected model was evaluated once on the held-out set to provide an independent estimate of performance. The development and held-out validation sets were separated at the patient level prior to any preprocessing or feature selection to prevent information leakage. Laboratory data spanned 18 monthly time bins across 10 myeloma-relevant analytes with 100% patient coverage; drug treatment records covered 74.4% of training set patients across three therapeutic classes.

For external validation, we used GSE24080 (UAMS-565), a publicly available microarray cohort of 507 multiple myeloma patients profiled on Affymetrix U133Plus2 microarrays with matched survival data (158 deaths, 31.1% event rate) [18, 26]. The higher event rate relative to CoMMpass reflects differences in treatment era, patient selection, and follow-up duration. Of the 5000 genes used in the development cohort, 4834 (96.7%) were matched by gene symbol in GSE24080; the remaining 166 unmatched genes were zero-filled. Critically, GSE24080 provides only baseline gene expression and five summary clinical variables (hemoglobin, creatinine, albumin, LDH and *β*_2_M); the longitudinal laboratory time series and drug treatment records that constitute two of the three modalities in the full teacher model are not available. Despite this modality mismatch, the external evaluation is designed to test whether a distilled student model— retaining only the DeepInsight image representation and baseline clinical features—can transfer meaningful prognostic discrimination to an independent cohort measured on a different platform.

### 2.2. Dynamic landmark prediction framework

The model operates in a dynamic landmark paradigm [17]: for a given patient and prediction time *t* ∈ {1, 2, …, 18} months, it observes all available data up to *t* and predicts residual survival risk over the subsequent one- to two-year horizon. This formulation is motivated by the clinical workflow in MM, where risk reassessment occurs at defined treatment milestones rather than at a single fixed time point. Unlike phase-specific prediction approaches that model events within a particular disease phase [8], the landmark framework produces continuously updated risk trajectories conditioned on the accumulated longitudinal records and thus enabling a single model to serve multiple clinical decision timepoints. This formulation ensures that predictions at each time point are based solely on information that would have been available in a real clinical setting.

A single architecture was used across all landmark times, with the current prediction time *t* encoded as a learned time embedding. Temporal integrity was enforced at the data level: the laboratory encoder received only time bins 1 through *t*, the treatment encoder received only pre-*t* drug records, and residual survival was defined as os days −*t* ×30. Only patients remaining at risk at time *t* contributed to evaluation at that landmark. Primary evaluation focused on *t* = 6 and *t* = 12 months.

### 2.3. Data modalities and preprocessing

Each modality was selected to capture a distinct layer of prognostic information in MM: the gene expression representation encodes baseline tumor biology and molecular subtype; the longitudinal laboratory data capture evolving disease burden and treatment response; and the drug history provides treatment context. This complementarity is a key design premise—the longitudinal modalities carry the most direct near-term prognostic signal (as confirmed by ablation), while the transcriptomic representation contributes a more stable but weaker baseline biological signal.

#### 2.3.1. Gene expression feature representation

From the full transcriptome, 5000 genes were selected by applying variance-based filtering on the training set, retaining only the genes with the highest expression variability across all patients. This subset captures the most informative transcriptomic variation while remaining computationally tractable for image construction. The selected genes were transformed into 96 × 96 single-channel images using DeepInsight [20]. By projecting high-dimensional transcriptomic data into a structured two-dimensional manifold, DeepInsight mitigates the curse of dimensionality and enables more efficient feature extraction compared to fully connected architectures. Gene coordinates in the image plane were determined by *t*-SNE (perplexity 30, cosine distance) fitted on the training set, mapping co-expressed genes to spatial proximity. *t*-SNE was selected for its ability to preserve local neighborhood structure, which is critical for capturing co-expression patterns relevant to functional gene modules. Expression values were min–max normalized on the training set and converted to pixel intensities via 1st/99th percentile clipping, such that each pixel represents the average normalized expression of genes mapped to that location. The DeepInsight transformation introduces an inductive bias aligned with biological organization: convolutional filters can exploit local co-expression patterns and functional gene modules, in contrast to fully connected architectures that treat genes as independent features.

#### 2.3.2. Longitudinal laboratory measurements

Ten analytes were selected for their established prognostic relevance in MM, including direct components of the ISS/R-ISS staging systems: hemoglobin (HGB), creatinine (CREAT), calcium (CA), leukocytes (WBC), albumin (ALB), free light chain lambda (FLC-*λ*), free light chain kappa (FLC-*κ*), serum M-protein, LDH, and *β*_2_M. Raw values were binned into monthly intervals, log-transformed where the distribution was right-skewed (FLC-*λ*, FLC-*κ*, LDH, *β*_2_M), and *z*-score normalized using training-set parameters. Missingness is an inherent feature of clinical laboratory data—sampling frequency varies by patient, institution, and clinical indication. To explicitly model this irregularity rather than treating it as nuisance, we accompany the observed values with binary observation masks **m** ∈ {0, 1}, per-analyte time since last observation (TSLO), and binary any-observation flags. This design allows the model to explicitly distinguish between true absence of signal and missing observations, a critical property in real-world clinical data.

#### 2.3.3. Treatment history

Drug exposure was encoded on the same monthly grid using three class-level binary indicators: bortezomib, carfilzomib, and immunomodulatory drugs (IMiDs). Each indicator takes value 1 if the patient received any agent within that drug class during the corresponding monthly interval and 0 otherwise; dosage information was not used, as the CoMMpass dataset records drug administration at the regimen level without standardized dose fields. Only pre-*t* records were visible at prediction time *t* to avoid data leakage.

### 2.4. Model architecture

The architecture follows a late-fusion multimodal design with a Cox proportional hazards (PH) prediction head (Figure 1B). Each modality is first encoded independently before being combined through a gated fusion mechanism that dynamically weights their contributions.

#### 2.4.1. Gene expression encoder

The 96 × 96 single-channel DeepInsight image is processed by a lightweight CNN comprising five convolutional blocks (each: 3 × 3 convolution, batch normalization, ReLU, 2×2 max-pooling), followed by adaptive average pooling and a fully connected projection layer with dropout rate 0.3. The encoder is fully trainable from the start, with its learning rate scaled by a factor of 0.1 relative to the base rate. The encoder contains ∼1.0M trainable parameters.

#### 2.4.2. Laboratory data encoder

A dual-stream Transformer architecture separately processes clinical measurements and observation profiles. The value stream takes as input, at each time step *j* ≤ *t*, the concatenation of observed lab values, binary masks, per-analyte TSLO, and any-observation flags, yielding a feature vector projected to dimension *d* = 128. Learnable positional embeddings are added, and the sequence is processed by a two-layer Transformer encoder (4 attention heads, feedforward dimension 256). A parallel missingness stream independently encodes observation patterns (TSLO, masks, any-observation flags) with a higher dropout to prevent the model from overfitting on missingness patterns. The two output streams are combined through a gating mechanism. Causal attention masks restrict each token to bins within [1, *t*].

#### 2.4.3. Drug encoder

A Transformer encoder (2 layers, 4 heads, dimension 64) processes the three-class drug usage grid with analogous TSLO features.

#### 2.4.4. Gated fusion

Modality embeddings are projected to a shared dimension (*d* = 128) and layer-normalized. A gate network—a two-layer MLP (hidden dimension 64, dropout 0.30) conditioned on **e**(*t*) and per-modality TSLO summaries—produces softmax-normalized weights. The fused representation is the weighted sum of projected modality embeddings. To mitigate modality collapse [16], two regularization mechanisms are applied: (i) auxiliary per-modality Cox PH heads that provide gradient supervision to each encoder (image head weight *λ*_img_ = 0.15; tabular head weight *λ*_tab_ = 0.10), and (ii) deploy-aware modality dropout (probability *p*_drop_ = 0.30) that jointly zeroes all clinical embeddings (laboratory data and drugs) during training to simulate external deployment scenarios where only the gene expression modality is available; the image representation modality is never dropped. This mechanism enables the model to down-weight modalities with sparse or outdated information, improving robustness to irregular clinical data.

#### 2.4.5. Prediction head and loss function

The fused representation is passed to the Cox PH head outputting a scalar log-hazard ratio *h*_*i*_ for each patient *i*. The model is trained by minimizing the negative Cox partial log-likelihood:

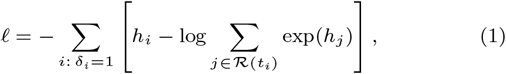

where *δ*_*i*_ denotes the event indicator and ℛ(*t*_*i*_) the risk set at time *t*_*i*_. The total training objective combines the fusion-level loss with auxiliary modality losses:

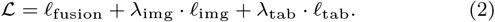

The complete teacher model contains ∼1.6M parameters. The Cox formulation was chosen for its robustness to censoring and its compatibility with dynamic risk prediction settings.

### 2.5. Training strategy

The training procedure was designed around two constraints specific to dynamic multimodal survival prediction in a small clinical cohort. The first constraint is sample efficiency: the development set contains only 624 patients with a 19.9% event rate, yet the model must learn shared representation across 18 landmark months, three heterogeneous modalities, and a gated fusion mechanism—a parameter-to-event ratio that is far less favorable than in imaging or NLP tasks of comparable architectural complexity. The second constraint is non-uniform event density across landmarks: early time points contribute more events and more patients to the risk set, while later landmarks are sparser, creating a systematic bias toward early-*t* patterns if training samples are drawn uniformly.

To address the first constraint, training operates at the patient level rather than the (patient, *t*) pair level: each patient contributes exactly *K* = 3 landmark samples per epoch, preventing over-representation of patients with longer follow-up and ensuring that the model sees every patient a controlled number of times. This design stabilizes gradient estimates and reduces variance in the Cox partial likelihood, which is particularly sensitive to risk-set composition when events are scarce. To address the second constraint, the landmark month *t* for each sample is drawn from a progressively broadened landmark sampling that initially concentrates on event-rich early landmarks— where gradient signal is strongest—and progressively broadens toward uniform coverage, ensuring that later landmarks receive adequate representation as the model matures. Together, these two mechanisms define a sampling contract: every patient is seen equally often, and every landmark receives attention proportional to its current learning utility. Together, these mechanisms ensure balanced exposure across patients and time points while maintaining stable gradient estimates.

Five-fold stratified cross-validation (stratified by event status) was performed within the 624-patient development set. Optimization used AdamW with linear warmup, plateau-based learning rate reduction, and gradient clipping—a conservative schedule chosen to avoid the sharp loss-landscape transitions that arise when Cox partial likelihood batches contain few events. Early stopping monitored exponentially smoothed validation tdAUC_1yr_ averaged across primary landmarks, with a minimum epoch gate to allow the curriculum to complete its transition before checkpointing begins. This curriculum stabilizes training by prioritizing informative early signals before incorporating more challenging later-stage patterns. Image augmentation was restricted to Gaussian noise and random erasing. Training convergence for the selected fold is shown in Supplementary Figure S1; both the Cox partial likelihood loss and validation Brier score decreased steadily. Full hyperparameter specifications—learning rates, batch sizes, scheduler parameters, and augmentation settings—are provided in Supplementary Methods.

### 2.6. Knowledge distillation and external deployment

External cohorts rarely provide the complete set of longitudinal modalities available in CoMMpass. To enable deployment on such cohorts, the multimodal teacher was distilled into a compact student model [22]. The student (∼1.0M parameters) receives only the DeepInsight image and five baseline clinical features: HGB, CREAT, ALB, LDH, and *β*_2_M—selected for their availability in the target external cohort (GSE24080) and their prognostic roles in ISS/R-ISS staging [3, 4, 5]. This reduced-input design represents a deployment-oriented compromise: the teacher’s full multimodal knowledge is compressed into a model that operates in settings where longitudinal data are unavailable. The student image encoder was initialized from teacher weights. Training used a composite objective:

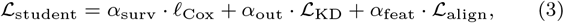

where 𝓁_Cox_ is the standard Cox partial log-likelihood (*α*_surv_ = 0.3), ℒ_KD_ is output-level distillation matching teacher and student log-hazard predictions (*α*_out_ = 0.5), and ℒ_align_ is feature-level cosine similarity (*α*_feat_ = 0.2). The best checkpoint was selected at epoch 86 (EMA-smoothed tdAUC_1yr_ = 0.831). External evaluation on GSE24080 (*n* = 507) proceeded without retraining; the 166 unmatched genes (of 5000) were zero-filled. This design enables practical deployment in settings where longitudinal data are unavailable while preserving key predictive signals learned by the full multimodal model.

### 2.7. Interpretability

Model interpretability was assessed at three levels: (1) Integrated Gradients [27] on the image representation branch, with pixel attributions back-mapped to genes via pyDeepInsight coordinates and linked to pathways through pre-ranked GSEA and ORA against MSigDB Hallmark gene sets [28]; (2) gradient-based temporal attribution of laboratory values across the 18 monthly time bins; and (3) drug counterfactual analysis estimating the change in predicted risk upon zeroing each treatment modality. These analyses provide complementary insights into both molecular and temporal drivers of risk predictions.

### 2.8. Evaluation metrics and benchmarks

Discrimination was assessed by Harrell’s concordance index (C-index) [29] and time-dependent AUC at 1-year and 2-year prediction horizons (tdAUC_1yr_, tdAUC_2yr_) computed via inverse probability of censoring weighting [30]. For risk stratification analysis, predicted risk scores were dichotomized into high- and low-risk groups using log-rank-optimized thresholds derived exclusively from the training set at each landmark; these thresholds were then applied without modification to the held-out validation and external cohorts. Kaplan– Meier survival curves were compared using the log-rank test, and hazard ratios were estimated from Cox regression on the dichotomized groups. Benchmark methods spanned three categories reflecting distinct modelling paradigms: (i) classical penalized survival regression (elastic net Cox), (ii) ensemble-based survival learning (random survival forest (RSF) [31], gradient-boosted survival analysis (GBSA)), and (iii) deep survival networks (DeepSurv [9], Cox-MLP). Each baseline was trained on three feature sets—clinical features, omics features (with PCA-50), and full concatenation— using matched CV splits, to separate the contribution of the learning algorithm from the input modality. This setup allows fair comparison across modelling paradigms while isolating the contribution of input modalities.

## 3. Results

### 3.1. Cross-validation performance and landmark-wise evaluation

The proposed multimodal framework achieved strong and consistent performance across all evaluation settings, demonstrating robust survival discrimination. Five-fold cross-validation on the development set (*n* = 624) yielded a mean C-index of 0.773 ± 0.024, tdAUC_1yr_ of 0.789 ± 0.021, and tdAUC_2yr_ of 0.798 ± 0.034 (per-fold results in Supplementary Table S1). Performance was stable across folds (C-index range 0.733–0.806). This low variability indicates that the model learns stable representation despite the limited sample size and event rate. On the held-out validation set (*n* = 128), the best model achieved C-indices of 0.793 at *t* = 6 months (*n* = 111, 17 events) and 0.819 at *t* = 12 months (*n* = 99, 10 events), with corresponding tdAUC_1yr_ of 0.813 and 0.818 (Supplementary Table S2).

Landmark-wise evaluation across prediction months *t* = 1 through 18 is shown in Figure 2. Both C-index and tdAUC_1yr_ remained in the 0.73–0.90 range, with discrimination increasing at later landmarks as more longitudinal information accumulated. This trend is consistent with expectations, as accumulating longitudinal information provides increasingly informative signals for risk estimation. It indicates that the model extracts meaningful prognostic signals across the full range of observation windows, not only from a single favorable landmark. Sample sizes decreased from *n* = 126 at *t* = 1 to *n* = 87 at *t* = 18.

**Figure 2.**
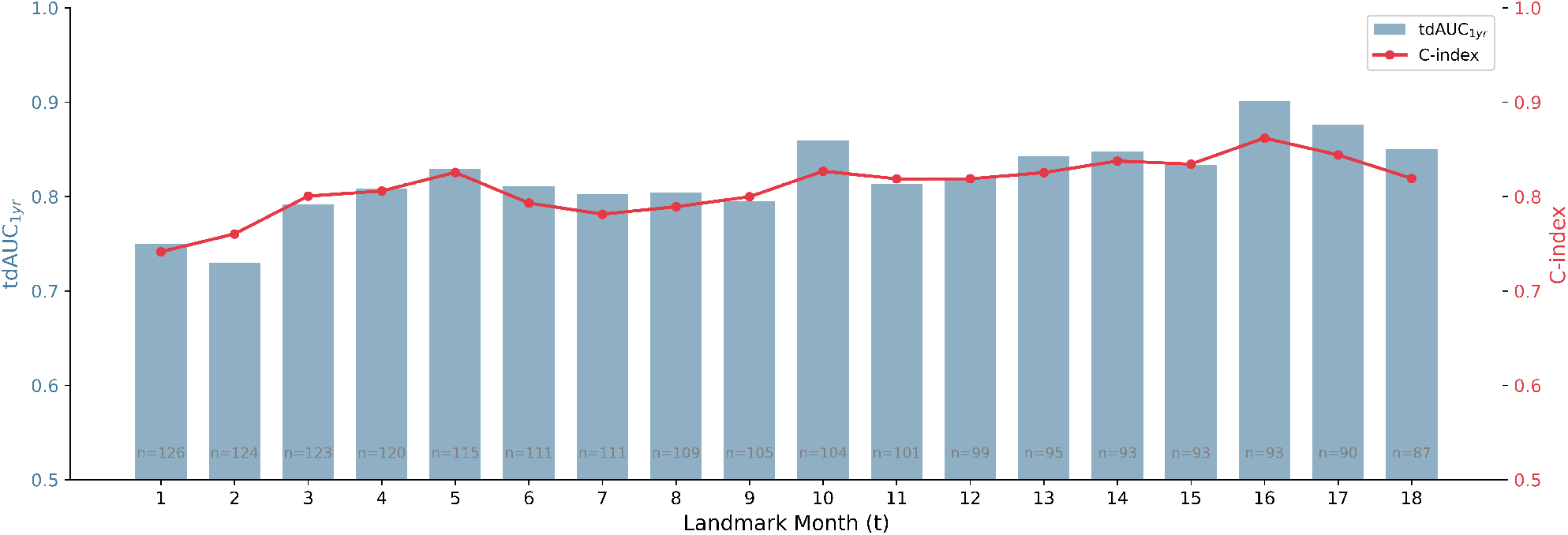
Landmark-wise evaluation across prediction months t = 1–18 on the held-out validation set (best fold). Bars: tdAUC_1yr_; line: C-index. Sample sizes annotated at each landmark.

### 3.2. Benchmark comparison

The proposed model consistently outperformed all baseline methods across all feature configurations and evaluation metrics. Figure 3 presents the best performance in the benchmark comparison on the held-out validation set; cross-fold stability is reported in Table 1 and Supplementary Figure S2. The proposed model (C-index 0.806, tdAUC_1yr_ 0.815, tdAUC_2yr_ 0.827) outperformed all baselines on the held-out set (Supplementary Table S3). Among methods receiving the full concatenated feature set, DeepSurv achieved C-index 0.733, Cox-MLP (a Cox model with multilayer perceptron feature extractor) 0.699, and elastic net 0.658. Among clinical-only models, RSF attained C-index 0.707 and GBSA 0.684. Omics-only models performed poorly (C-index 0.640–0.672).

**Table 1.**
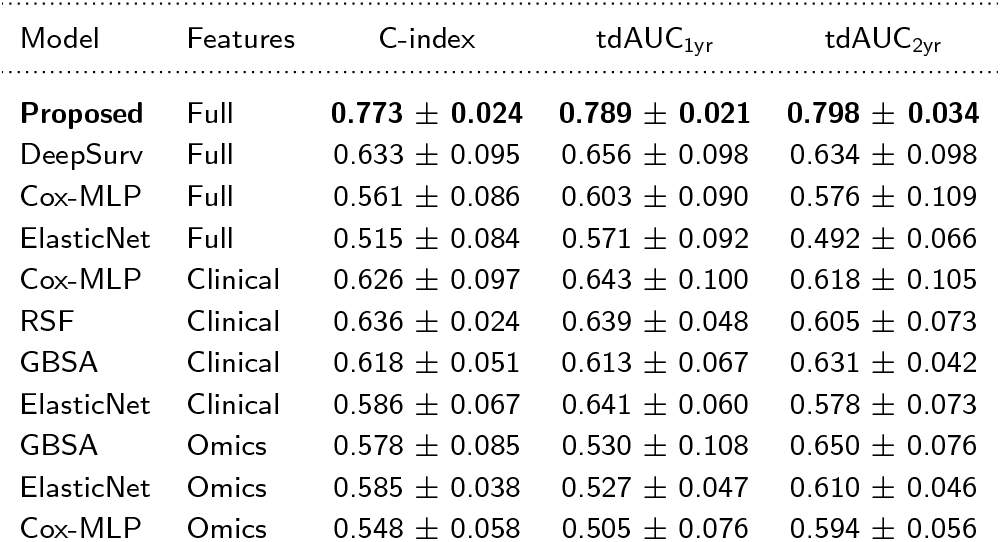
Benchmark comparison (five-fold CV, mean SD across folds). All models were trained and evaluated independently on identical fold splits generated by a single stratified partitioning. Best performance in bold.

| Model | Features | C-index | tdAUC<sub>1yr</sub> | tdAUC<sub>2yr</sub> |
| --- | --- | --- | --- | --- |
| **Proposed** | Full | **0.773 ± 0.024** | **0.789 ± 0.021** | **0.798 ± 0.034** |
| DeepSurv | Full | 0.633 ± 0.095 | 0.656 ± 0.098 | 0.634 ± 0.098 |
| Cox-MLP | Full | 0.561 ± 0.086 | 0.603 ± 0.090 | 0.576 ± 0.109 |
| ElasticNet | Full | 0.515 ± 0.084 | 0.571 ± 0.092 | 0.492 ± 0.066 |
| Cox-MLP | Clinical | 0.626 ± 0.097 | 0.643 ± 0.100 | 0.618 ± 0.105 |
| RSF | Clinical | 0.636 ± 0.024 | 0.639 ± 0.048 | 0.605 ± 0.073 |
| GBSA | Clinical | 0.618 ± 0.051 | 0.613 ± 0.067 | 0.631 ± 0.042 |
| ElasticNet | Clinical | 0.586 ± 0.067 | 0.641 ± 0.060 | 0.578 ± 0.073 |
| GBSA | Omics | 0.578 ± 0.085 | 0.530 ± 0.108 | 0.650 ± 0.076 |
| ElasticNet | Omics | 0.585 ± 0.038 | 0.527 ± 0.047 | 0.610 ± 0.046 |
| Cox-MLP | Omics | 0.548 ± 0.058 | 0.505 ± 0.076 | 0.594 ± 0.056 |

**Figure 3.**
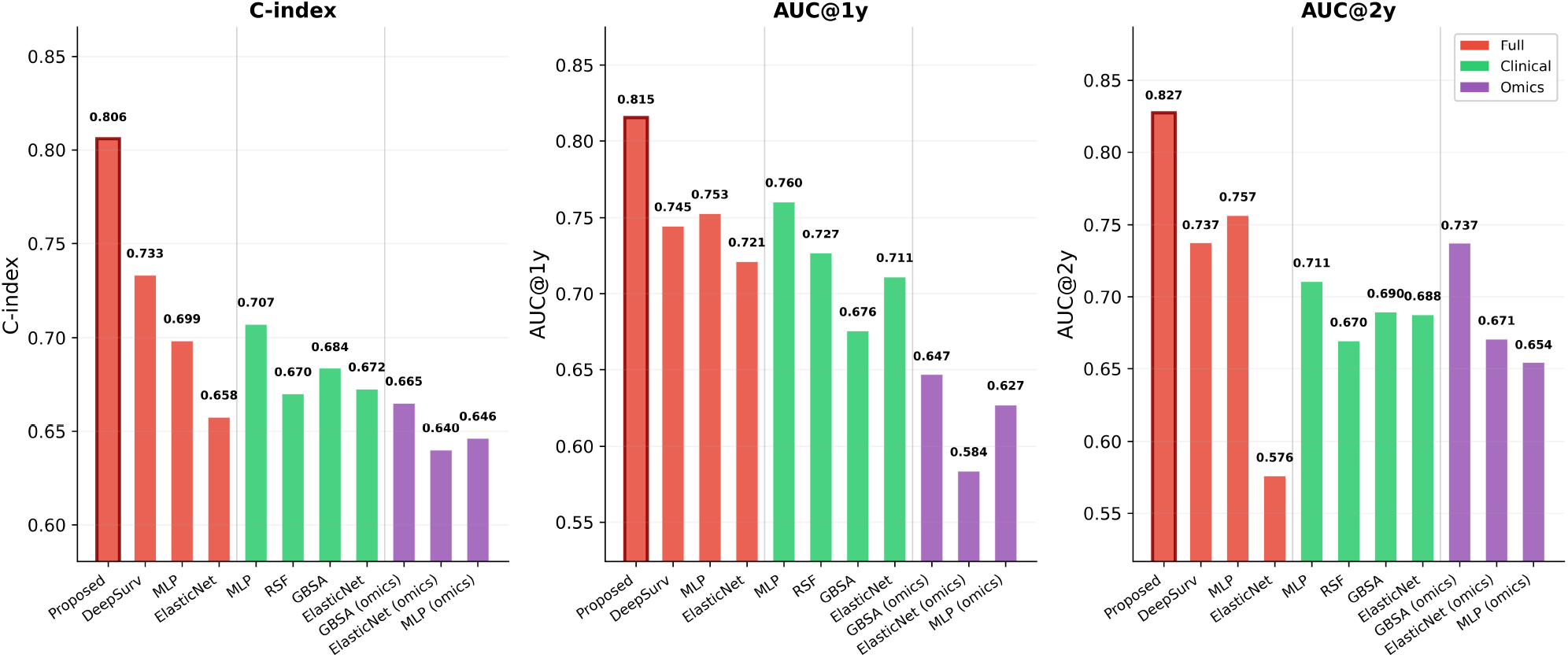
Benchmark comparison on the held-out validation set (best fold). C-index, tdAUC_1yr_, and tdAUC_2yr_ for the proposed model and all baselines, grouped by input modality.

The five-fold CV aggregate confirmed this ordering. The proposed model achieved a mean C-index of 0.773 ± 0.024, compared with the next-best RSF at 0.636 ± 0.024 and DeepSurv at 0.633 ± 0.095.

The proposed model exhibits lower cross-fold variability (SD 0.024) than most deep learning baselines (DeepSurv SD 0.095; Cox-MLP SD 0.086–0.097), suggesting more stable learning.

### 3.3. Overall survival risk stratification

Kaplan–Meier analysis on the held-out validation set showed significant survival risk-group separation at both primary landmarks (Figure 4). At *t* = 6 months, the high-risk group (*n* = 21) had lower survival than the low-risk group (*n* = 90), with a hazard ratio (HR) of 3.46 (log-rank *p <* 0.001). At *t* = 12 months, separation was more pronounced (HR = 3.93, *p <* 0.001; high-risk *n* = 18, low-risk *n* = 81). The larger HR at later landmarks suggests improved risk separation as more longitudinal information becomes available. These results demonstrate that the model’s predictions support clinically meaningful patient stratification. In addition, Kaplan– Meier–based grouped calibration at the 12-month landmark indicated that predicted and observed survival probabilities were broadly consistent, with mean absolute errors of 0.118 for the 1-year horizon and 0.088 for the 2-year horizon (Supplementary Figure S3).

**Figure 4.**
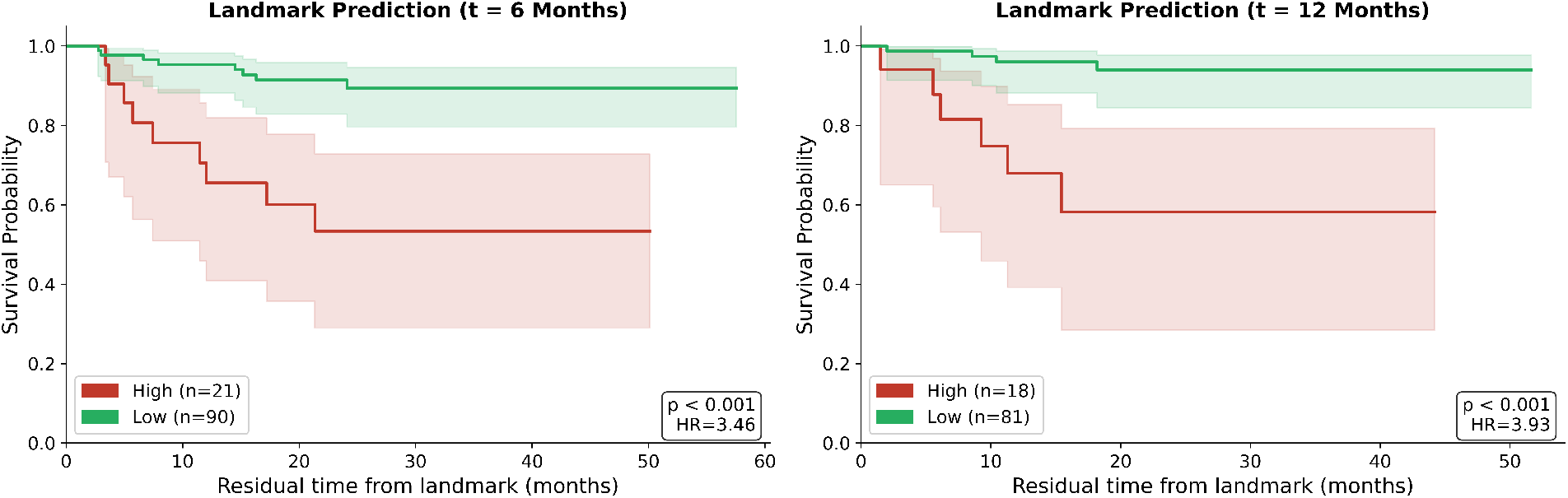
Kaplan–Meier survival curves stratified by model-predicted overall survival risk at landmarks *t* = 6 months (left) and *t* = 12 months (right) on the held-out validation set. Shaded bands: 95% confidence intervals. Hazard ratios and log-rank *p*-values annotated.

### 3.4. Cross-platform external validation

To evaluate cross-platform generalization and external validity, we evaluated a distilled version of the model on an independent external cohort without retraining. The distilled student model was applied to GSE24080 (*n* = 507, 158 deaths). It is important to note that this evaluation tests not the full teacher model’s generalizability but the viability of a reduced-input deployment path: whether a distilled model retaining only baseline features can transfer meaningful prognostic discrimination to an independent cohort measured on a different platform.

For external validation, we used the fold-specific checkpoint with the best validation performance for each benchmarked model, as determined on the held-out CoMMpass validation set. At the baseline landmark (*t* = 1 month), the student achieved C-index 0.672 and tdAUC_1yr_ 0.740, outperforming all transferred baselines: DeepSurv (C-index 0.645), Cox-MLP clinical (0.628), RSF clinical (0.610), and GBSA clinical (0.592) (Figure 5A; Supplementary Table S4). Multi-landmark evaluation showed stable performance from *t* = 1 through 12 months (C-index 0.649–0.672, tdAUC_1yr_ 0.710– 0.755; Supplementary Table S5). Figure 5B shows representative Kaplan–Meier stratification at landmarks *t* = 3, 6, 9 months, all demonstrating significant risk-group separation (log-rank *p <* 10^−10^); complete results are reported in Supplementary Table S5.

**Figure 5.**
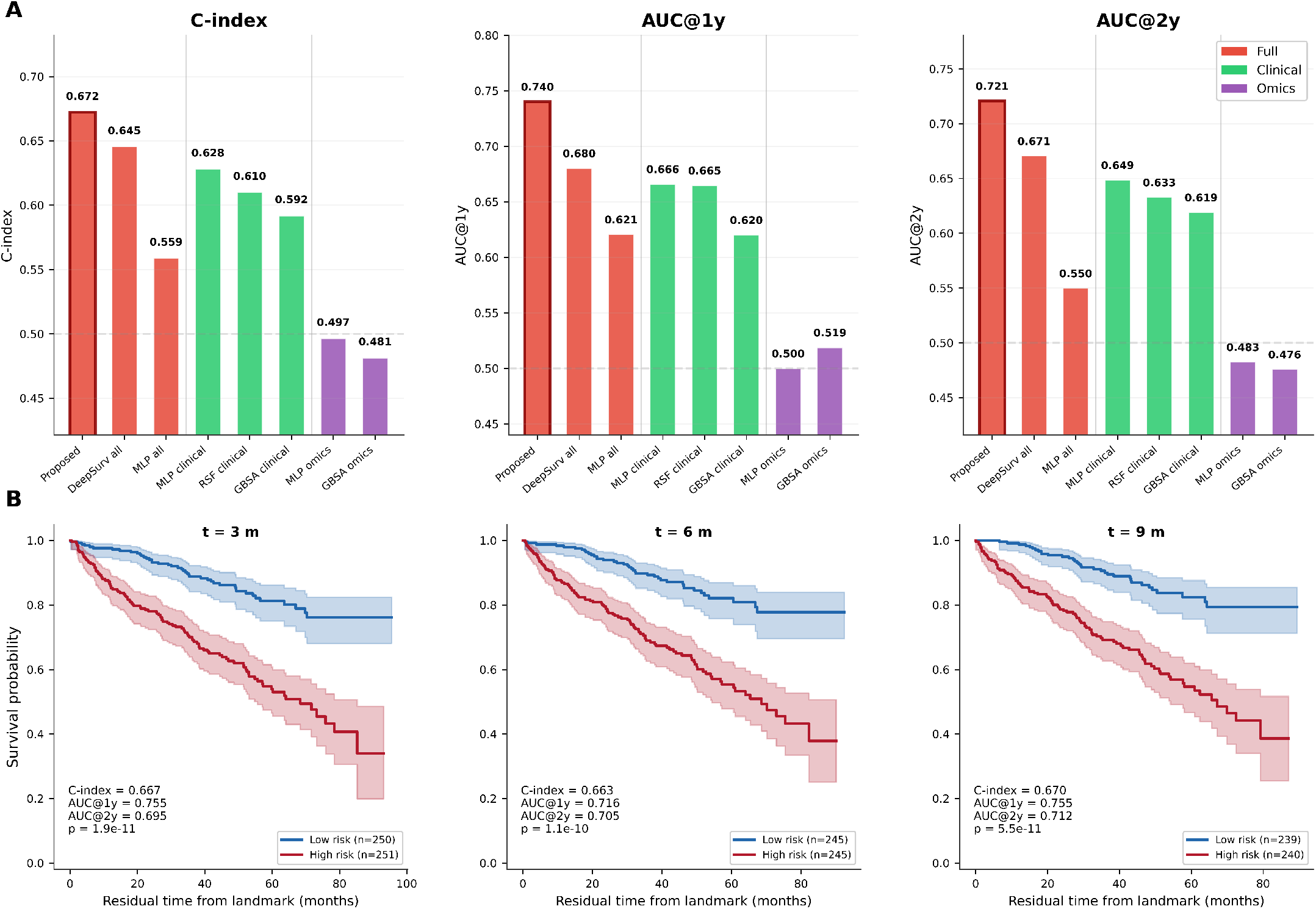
External validation on GSE24080 (*n* = 507). **(A)** Benchmark comparison at *t* = 1 month. **(B)** Kaplan–Meier survival curves at representative landmarks *t* = 3, 6, 9 months. C-index, log-rank *p*-values, and tdAUC metrics annotated. Complete multi-landmark results are provided in Supplementary Table S5.

This demonstrates that the learned representation retains predictive value across cohorts and measurement platforms.

Modality ablation on the external cohort (Supplementary Table S6; Supplementary Figure S4) confirmed that the combined image representation and clinical features (C-index 0.672) outperformed clinical features alone (0.659) and image representation alone (0.595). Overall, these results support the feasibility of deploying a reduced-input model that preserves key multimodal insights in limited data clinical settings.

## Discussion

### 4.1. Performance and dynamic prediction

Here we developed a multimodal deep learning framework for dynamic survival prediction in MM that integrates gene expression image representation, longitudinal laboratory tests, and treatment context through gated fusion. The model produces updated risk estimates from any observation window within the first 18 months of diagnosis while enforcing temporal integrity. Together, these features enable clinically aligned, time-adaptive risk prediction that extends beyond conventional static prognostic models.

The internal C-index of 0.806 is comparable to previously reported values on the CoMMpass cohort: Mosquera Orgueira et al. [7] achieved 0.780 with random forests combining clinical and RNA-seq features, and Hussain et al. [8] reported 0.70–0.75 with SCOPE. Direct comparison across studies is confounded by differences in feature sets, cohort definitions, and evaluation protocols. Beyond such differences in datasets, key distinction lies in the modelling paradigm: prior approaches relied on static or phase-specific features, whereas the proposed framework incorporates time-evolving clinical information within a unified dynamic model. This design enables the model to capture treatment response dynamics and evolving risk profiles that fixed-time-point architectures cannot access.

### 4.2. Modality ablation

Single modality ablation (Table 2; Supplementary Figure S5; Supplementary Figure S6) quantified each data source’s contribution and revealed a clinically interpretable hierarchy. Longitudinal laboratories were the strongest individual modality (C-index 0.693 ± 0.021, tdAUC_1yr_ 0.739 ± 0.026), consistent with the centrality of *β*_2_M, albumin, and LDH to ISS/R-ISS staging [3, 4, 5] and reflecting the fact that longitudinal trajectories capture the most direct near-term disease-state signal. Drug history alone reached C-index 0.628 ± 0.045; the relatively weaker standalone performance is expected, as treatment intensity correlates with disease severity (indication bias), making drug features more informative as a complement to clinical data than as an independent predictor. The DeepInsight image representation branch (0.624 ± 0.029) outperformed a flat expression MLP model on the same 5000 input genes (0.596 ± 0.044; ΔC ≈ 0.028), suggesting that the spatial encoding of co-expression structure provides a useful inductive bias. While modest in absolute terms, such improvements are meaningful in survival modelling, where gains of 0.02–0.03 in C-index are typically considered non-trivial [29]. This suggests that spatial encoding enables the model to capture co-expression patterns that are not readily accessible to vector-based models.

**Table 2.** Modality ablation (five-fold CV, mean ± SD).

| Configuration | C-index | tdAUC<sub>1yr</sub> | $\Delta C$ |
| --- | --- | --- | --- |
| Full (Proposed) | 0.773±0.024 | 0.789±0.021 | — |
| Lab only | 0.693±0.021 | 0.739±0.026 | −0.080 |
| Drug only | 0.628±0.045 | 0.625±0.060 | −0.145 |
| DeepInsight image | 0.624±0.029 | 0.590±0.030 | −0.149 |
| Expression MLP | 0.596±0.044 | 0.575±0.042 | −0.177 |

The full multimodal model improved over the best single modality by ΔC = +0.080 (10.4% relative), indicating that the three data sources carry complementary prognostic information that the gated fusion mechanism successfully combines.

### 4.3. Molecular and clinical interpretability

The model’s predictions were further examined for biological and clinical interpretability. Integrated Gradients on the gene expression image representation branch produced pixel-level attribution maps that were reverse-mapped to individual genes via pyDeepInsight coordinates (Figure 6A–C). Among risk-associated genes, UBE2Q1 had the highest mean Integrated Gradients attribution. UBE2Q1 encodes a ubiquitin-conjugating enzyme within the ubiquitin-proteasome system (UPS), a pathway central to myeloma biology and to the therapeutic activity of proteasome inhibitors [32, 33]. Other high-ranking risk genes included ADAR, whose overexpression in MM has been linked to aberrant A-to-I hyper-editing, oncogenicity, and poor prognosis [34]; P4HB, a protein disulfide isomerase involved in endoplasmic reticulum protein folding, consistent with the marked dependence of myeloma plasma cells on ER homeostasis and their vulnerability to unfolded protein response disruption [35]; and PRPF8, a core spliceosome component, consistent with emerging evidence that RNA splicing is dysregulated in multiple myeloma and that the spliceosome may represent a therapeutic vulnerability in this disease [36]. Among protective genes, SPEN (a transcriptional repressor in the Notch signaling pathway) and RNF169 (a DNA damage response factor) were prominent. Representative attribution maps for high-risk and low-risk patients are provided in Supplementary Figure S7, illustrating the spatial concentration of high-magnitude attributions in the high-risk group relative to the diffuse patterns in the low-risk group.

**Figure 6.**
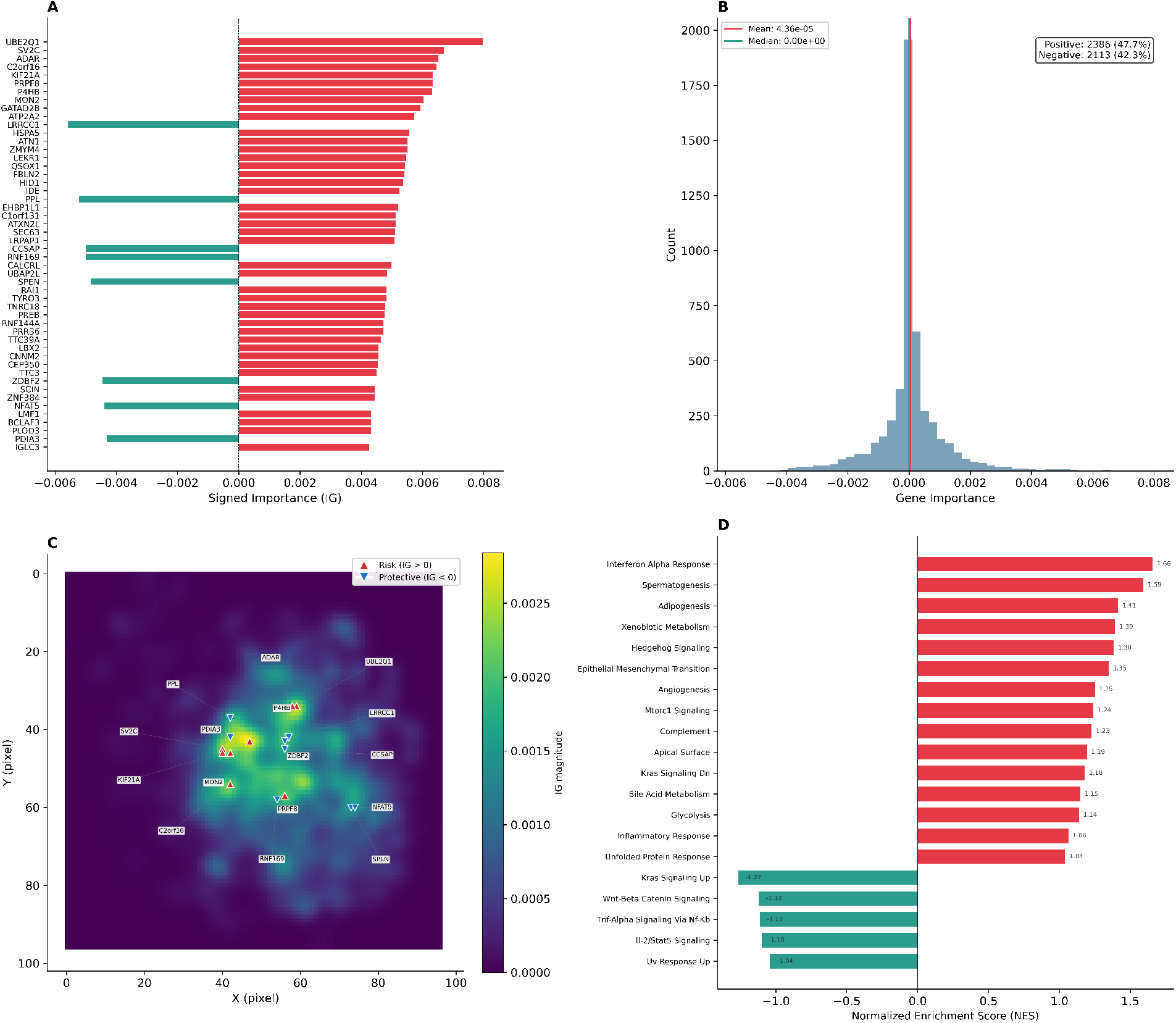
Gene-level interpretability. **(A)** Top genes ranked by signed Integrated Gradients attribution. **(B)** Distribution of gene-level importance scores. **(C)** Spatial importance map on DeepInsight coordinates. **(D)** Pathway enrichment via pre-ranked GSEA on MSigDB Hallmark gene sets.

Pre-ranked GSEA on MSigDB Hallmark gene sets (Figure 6D) identified risk-associated enrichment in Interferon Alpha Response, consistent with evidence that type I interferon signaling can promote myeloma-associated immunosuppression and disease progression [37]. ORA on both risk-increasing and protective gene sets (Supplementary Figures S8 and S9) also identified Unfolded Protein Response and mTORC1 Signaling among risk-enriched terms. These associations support biological plausibility but do not establish causal relationships; the model captures correlative transcriptomic patterns that align with known myeloma biology rather than identifying therapeutic targets.

Temporal attribution of laboratory values was computed by backpropagating the model’s log-hazard output through the laboratory encoder and recording the signed gradient with respect to each observed analyte value at each monthly time bin. For a given analyte at a certain time bin, a positive gradient indicates that increasing the observed value at that time point would raise the predicted mortality risk, while a negative gradient indicates a protective association. The resulting heatmap (Figure 7) revealed consistent direction structure across analytes: FLC-*κ*, FLC-*λ*, LDH, and *β*_2_M exhibited positive gradients throughout the observation window, confirming their established associations with disease burden. Albumin showed the strongest negative (protective) gradients, followed by hemoglobin—consistent with the known prognostic roles of these markers in ISS/R-ISS staging [3, 4, 5].

**Figure 7.**
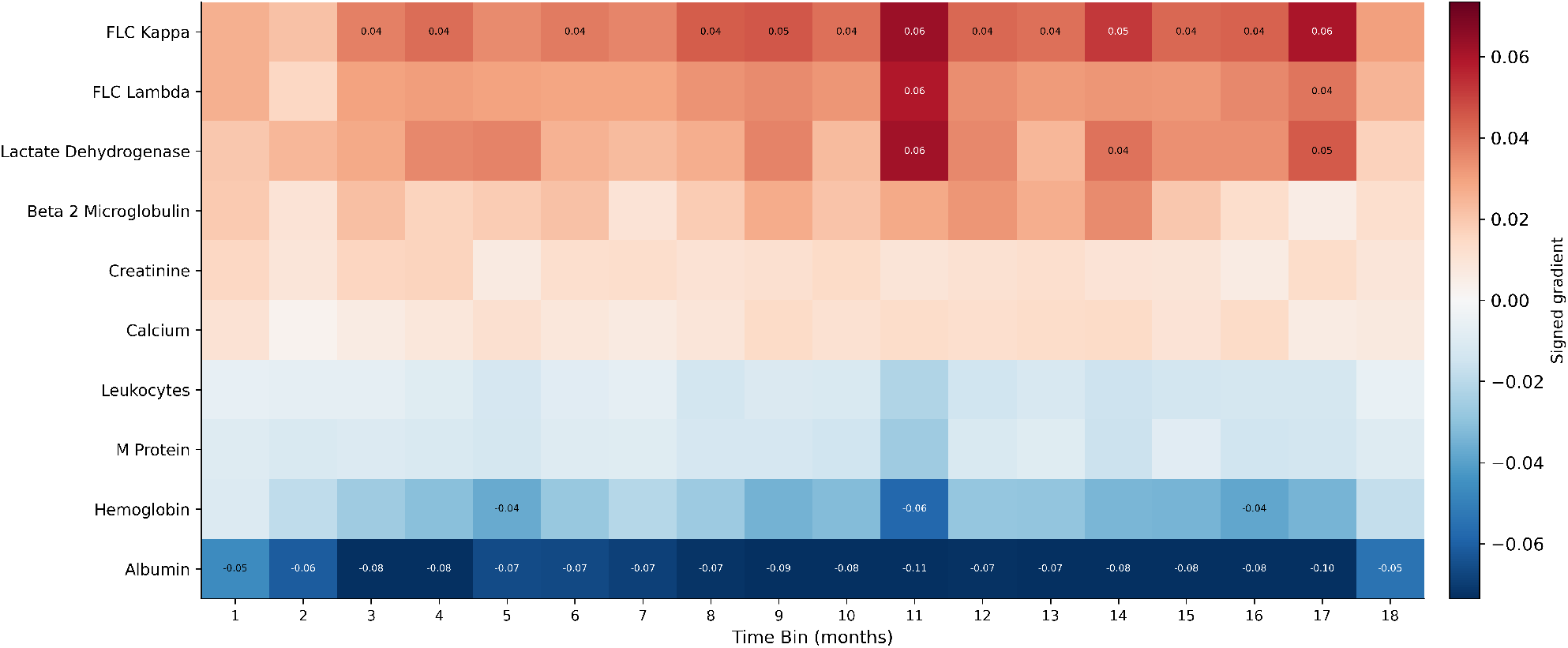
Laboratory temporal attribution heatmap. Each cell shows the mean signed gradient *∂*(log-hazard)*/∂*(analyte value) at the corresponding monthly time bin. Positive values (red) indicate that higher analyte values increase predicted mortality risk; negative values (blue) indicate a protective association. Analytes are sorted by mean gradient direction (risk-associated above, protective below).

To characterize treatment context, we examined drug utilization patterns stratified by model-predicted risk group (Supplementary Figure S10). Patients in the high-risk group (death rate 34%) received more intensive treatment than those in the low-risk group (death rate 1%): carfilzomib and IMiD usage rates were consistently higher across all monthly time bins, with the largest differences emerging after month 5. This pattern is consistent with clinical practice, where higher-risk patients are more likely to receive second-line proteasome inhibitors and combination regimens.

### 4.4. Limitations and future directions

This study has several limitations. The generalizability of the proposed framework across populations with different demographic profiles, treatment patterns, and follow-up structures remains to be determined. Although internal five-fold cross-validation supported stable performance within the CoMMpass cohort, external assessment on GSE24080 was conducted using a reduced-input student model because that cohort lacks longitudinal laboratory measurements and treatment histories. Consequently, the current external results provide limited evidence regarding cross-cohort generalizability of the full framework. Further evaluation in independent cohorts with harmonized multimodal data collection and longitudinal follow-up will be required to determine its utility in longitudinal clinical settings. In addition, the model was trained using the Cox partial likelihood, which relies on the proportional hazards assumption; this assumption may not hold in certain clinical subgroups, including patients with late-relapsing multiple myeloma. Future studies should therefore investigate alternative survival modeling objectives, including rank-based losses and discrete-time formulations, to assess whether they provide greater robustness under non-proportional hazard settings.

## 5. Conclusion

We developed a dynamic multimodal survival prediction framework for multiple myeloma that integrates gene expression, longitudinal laboratory trajectories, and treatment history through gated fusion.

The model produces updated risk estimates from clinically relevant time points while preserving temporal integrity. The framework achieved strong and consistent performance: five-fold cross-validation on the CoMMpass cohort yielded a C-index of 0.773 ± 0.024 and tdAUC_1yr_ of 0.789 ± 0.021. A distilled student model retained discriminative power on an independent external cohort (C-index 0.672, tdAUC_1yr_ 0.740), despite platform differences and restricted inputs. Interpretability analyses identified molecular and clinical patterns consistent with known myeloma biology, supporting the biological plausibility of the model’s predictions. Overall, these results reaffirm the potential of dynamic multimodal frameworks to improve prognostic modelling in longitudinal clinical settings such as multiple myeloma. Future work will focus on prospective validation with complete multimodal data through clinical collaboration to assess translational utility.

## Data availability

The primary development cohort was derived from the MMRF CoMMpass study [24, 25], available via the MMRF Researcher Gateway subject to data-access requirements. External validation data are publicly available from the NCBI Gene Expression Omnibus (GEO) under accession GSE24080 [26]. A reproducibility package containing preprocessed data, trained model checkpoints, and training outputs is archived on figshare (https://doi.org/10.6084/m9.figshare.31804762).

## Code availability

The source code has been modularized and made publicly available at the GitHub repository: https://github.com/shangruJia/multi_mm.

## Author contributions

SJ implemented the whole pipeline, evaluated the performance, and wrote the first draft and contributed to the subsequent versions of the manuscript. AL advised for the model and the evaluation, and contributed to the manuscript writeups. KAB checked the model, and helped in the manuscript writeup. AS and TT perceived, supervised, and contributed to the manuscript writeups. All authors read and approved the manuscript.

## Acknowledgments

The results presented in this study are based in part on data generated by the MMRF CoMMpass Study and on the publicly available GSE24080 dataset. We gratefully acknowledge the patients and their families, and the investigators and consortia who generated and shared these valuable datasets. Their contributions made possible the model development, benchmarking, biological interpretation, and external validation reported in this work.

