## Supplementary File for "Dynamic multimodal survival prediction in multiple myeloma integrating gene expression, longitudinal laboratories, and treatment history"

#### Supplementary Tables

**Table S1:** Performance of the five fold-specific models on the common held-out CoMMpass validation split ($n = 128$).

| Fold | C-index | tdAUC$_{1yr}$ | tdAUC$_{2yr}$ | tdAUC$_{1yr}$ $t$=6 | Status |
|---|---|---|---|---|---|
| 0 | 0.776 | 0.812 | 0.791 | 0.861 | Completed |
| 1 | 0.780 | 0.774 | 0.811 | 0.775 | Completed |
| 2 | 0.806 | 0.815 | 0.826 | 0.813 | **Selected** |
| 3 | 0.733 | 0.778 | 0.735 | 0.741 | Completed |
| 4 | 0.770 | 0.764 | 0.827 | 0.758 | Completed |
| Mean ± SD | 0.773±0.024 | 0.789±0.021 | 0.798±0.034 | 0.790±0.043 | — |

**Note.** Five fold-specific models trained on stratified partitions of the 624-patient development set were compared on a common held-out CoMMpass validation split ($n = 128$). Fold 2 was selected for downstream analyses based on the highest mean tdAUC$_{1yr}$ across the primary landmarks ($t = 6, 12$ months).

**Table S2:** Held-out validation performance at primary landmarks for the selected fold (fold 2, $n = 128$). Metrics were computed using inverse-probability-of-censoring weighting.

| Landmark | $n$ | Events (%) | C-index | tdAUC$_{1yr}$ | tdAUC$_{2yr}$ |
|---|---|---|---|---|---|
| $t = 6$ months | 111 | 17 (15.3%) | 0.793 | 0.813 | 0.816 |
| $t = 12$ months | 99 | 10 (10.1%) | 0.819 | 0.818 | 0.837 |

**Note.** Aggregate: tdAUC$_{1yr}$ avg = 0.815; tdAUC$_{2yr}$ avg = 0.826; C-index avg = 0.806.

**Table S3:** Benchmark comparison on the held-out validation set for the selected fold (fold 2, $n = 128$). All models were evaluated on matched data splits. Best value in each column is shown in bold.

| Model | Features | C-index | $\text{tdAUC}_{1\text{yr}}$ | $\text{tdAUC}_{2\text{yr}}$ |
|---|---|---|---|---|
| **Proposed** | Full | **0.806** | **0.815** | **0.827** |
| DeepSurv | Full | 0.733 | 0.745 | 0.737 |
| Cox-MLP | Full | 0.699 | 0.721 | 0.757 |
| ElasticNet | Full | 0.658 | 0.753 | 0.711 |
| Cox-MLP | Clinical | 0.707 | 0.760 | 0.711 |
| RSF | Clinical | 0.707 | 0.727 | 0.670 |
| GBSA | Clinical | 0.684 | 0.676 | 0.690 |
| ElasticNet | Clinical | 0.670 | 0.711 | 0.670 |
| GBSA | Omics | 0.672 | 0.647 | 0.688 |
| ElasticNet | Omics | 0.665 | 0.627 | 0.690 |
| Cox-MLP | Omics | 0.646 | 0.584 | 0.654 |

**Table S4:** External validation benchmark on the GSE24080 cohort at the baseline landmark ($t = 1$ month, $n = 507$, 158 events).

| Model | Features | C-index | $\text{tdAUC}_{1\text{yr}}$ | $\text{tdAUC}_{2\text{yr}}$ |
|---|---|---|---|---|
| **Proposed (Student)** | Image + Clinical | **0.672** | **0.740** | **0.721** |
| DeepSurv | All | 0.645 | 0.680 | 0.671 |
| Cox-MLP | Clinical | 0.628 | 0.666 | 0.649 |
| RSF | Clinical | 0.610 | 0.665 | 0.633 |
| GBSA | Clinical | 0.592 | 0.620 | 0.619 |
| Cox-MLP | All | 0.559 | 0.621 | 0.550 |
| Cox-MLP | Omics | 0.497 | 0.500 | 0.483 |
| GBSA | Omics | 0.481 | 0.519 | 0.476 |

**Note.** For external transfer, each benchmarked method was instantiated using the fold-specific model selected on the common held-out CoMMpass validation split. The evaluation was performed without any use of GSE24080 for model or threshold selection.

**Table S5:** Multi-landmark external validation on GSE24080 using the distilled student model. Performance at each landmark was evaluated independently; $n$ decreases at later landmarks as patients with shorter follow-up are excluded from the at-risk set.

| $t$ (months) | $n$ | C-index | $\text{tdAUC}_{1\text{yr}}$ | $\text{tdAUC}_{2\text{yr}}$ |
|---|---|---|---|---|
| 1 | 507 | 0.672 | 0.740 | 0.721 |
| 3 | 501 | 0.667 | 0.755 | 0.695 |
| 6 | 490 | 0.663 | 0.716 | 0.705 |
| 9 | 479 | 0.670 | 0.755 | 0.712 |
| 12 | 468 | 0.649 | 0.710 | 0.669 |

**Table S6:** Modality ablation for the distilled student model on GSE24080 ($t = 1$, $n = 507$). The full student model combines the DeepInsight image representation with five baseline clinical features (HGB, CREAT, ALB, LDH, $\beta_2$M).

| Configuration | C-index | $\text{tdAUC}_{1\text{yr}}$ | $\text{tdAUC}_{2\text{yr}}$ |
|---|---|---|---|
| Full (Image + Clinical) | **0.672** | **0.740** | **0.721** |
| Image only | 0.595 | 0.580 | 0.607 |
| Clinical only | 0.659 | 0.734 | 0.712 |

**Note.** $\Delta$C over clinical-only: +0.013; $\Delta$C of clinical over image-only: +0.064.

### Supplementary Figures

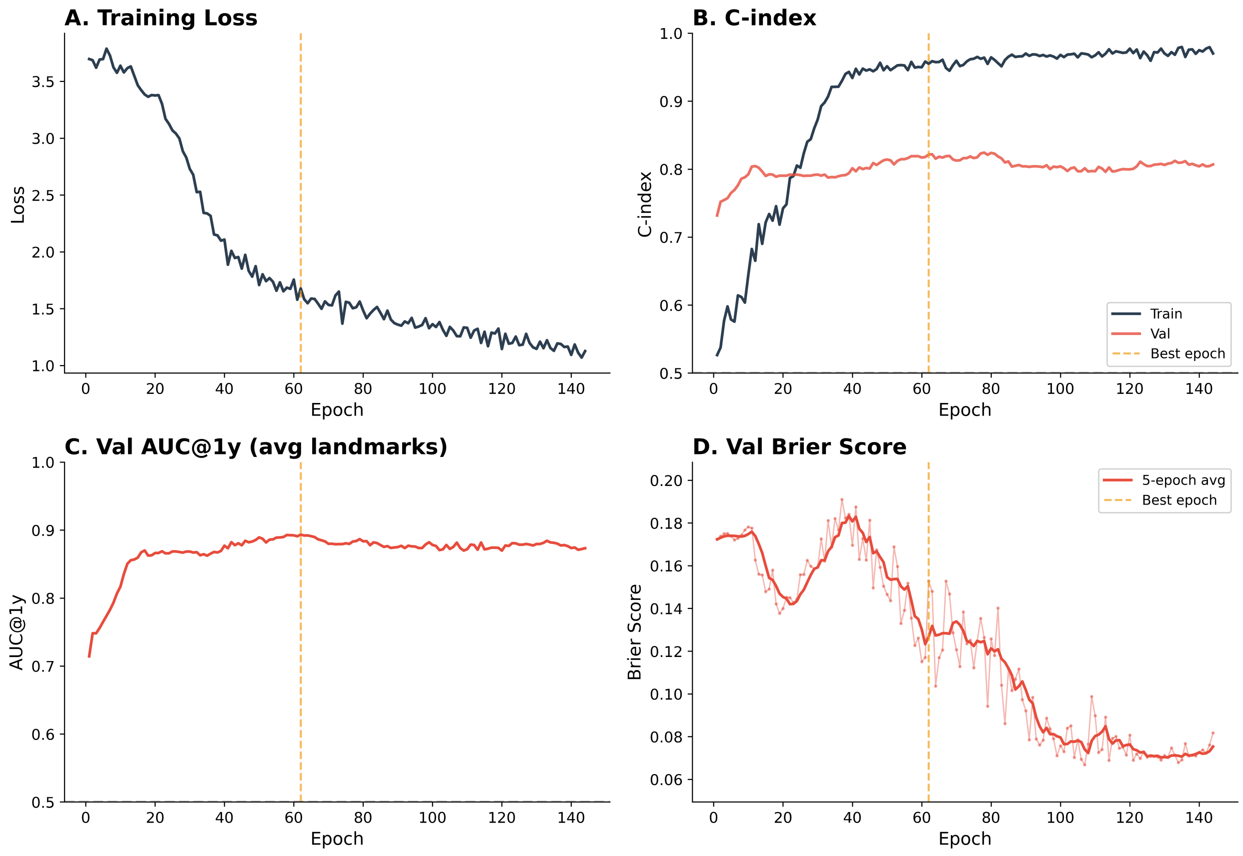


**Figure S1:** Training convergence for the selected fold (fold 2). **(A)** Negative Cox partial log-likelihood loss over training epochs. **(B)** Training and validation C-index trajectories; the vertical dashed line marks the best epoch (epoch 62) as determined by the EMA-smoothed $\text{tdAUC}_{1\text{yr}}$ monitor. **(C)** Validation $\text{tdAUC}_{1\text{yr}}$ averaged across primary landmarks ($t = 6, 12$ months), used as the early stopping criterion. **(D)** Validation Brier score (1-year horizon); the red line shows the 5-epoch rolling mean. Validation metrics were computed using model-weight exponential moving average (Polyak averaging, decay 0.998).

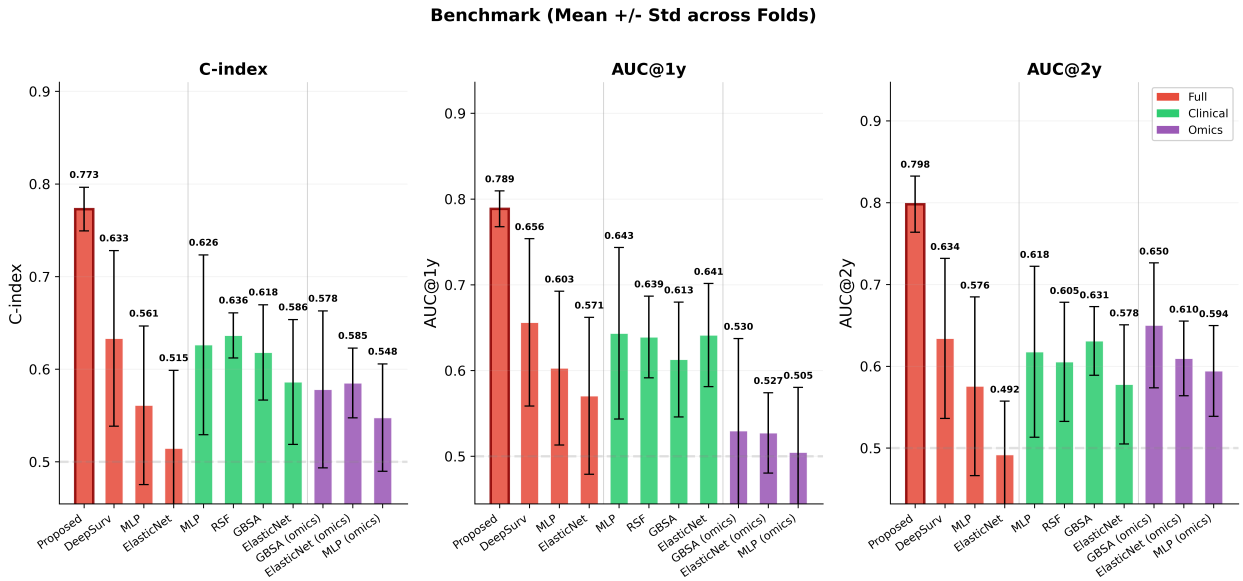


**Figure S2:** Benchmark comparison across five-fold cross-validation (mean ± SD). C-index, $\mathrm{tdAUC}_{1\mathrm{yr}}$, and $\mathrm{tdAUC}_{2\mathrm{yr}}$ are shown for the proposed model and all baselines, grouped by input feature set. Error bars represent one standard deviation over five folds computed on matched stratified splits. This figure complements main-text Table 1 (fold-level means) and Figure 3 (best-fold point estimates).

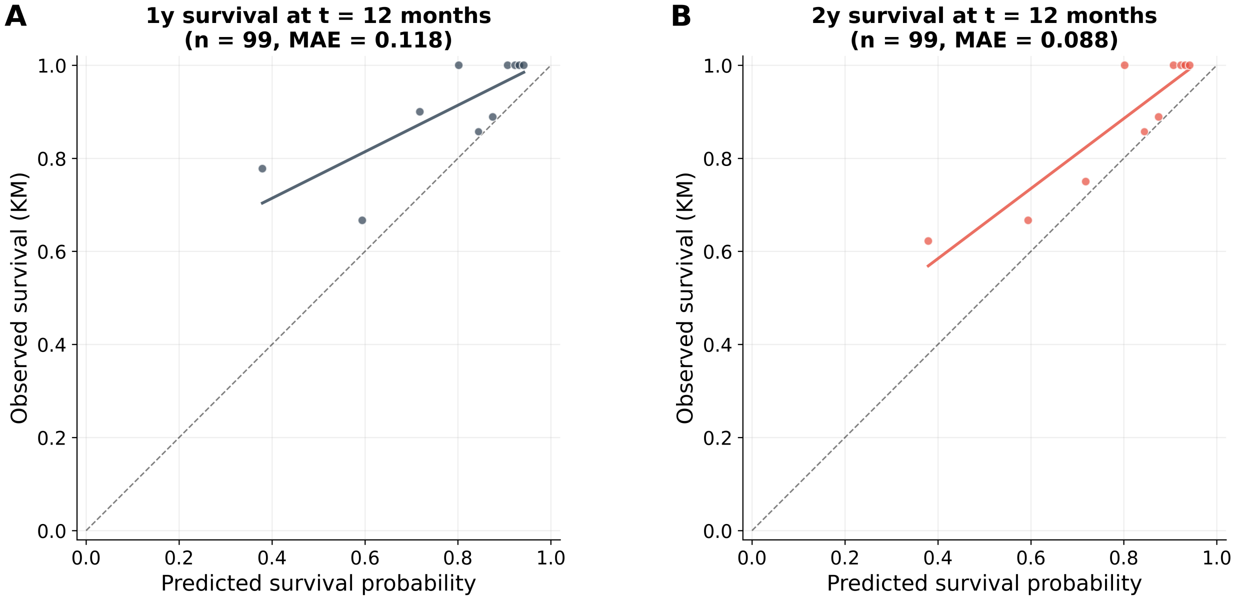


**Figure S3:** Calibration assessment at the $t = 12$ month landmark on the held-out validation set ($n = 99$). **(A)** Predicted versus Kaplan–Meier observed 1-year survival probability (mean absolute error, MAE = 0.118). **(B)** Predicted versus Kaplan–Meier observed 2-year survival probability (MAE = 0.088). Each point represents a patient-level predicted–observed pair obtained by stratifying predicted survival probabilities into quantile bins and computing the corresponding Kaplan–Meier estimate within each bin. The diagonal dashed line indicates perfect calibration; solid lines show locally weighted regression fits. The lower MAE at the 2-year horizon reflects the model's tendency toward conservative (higher) survival probability estimates that align more closely with observed outcomes over longer follow-up.

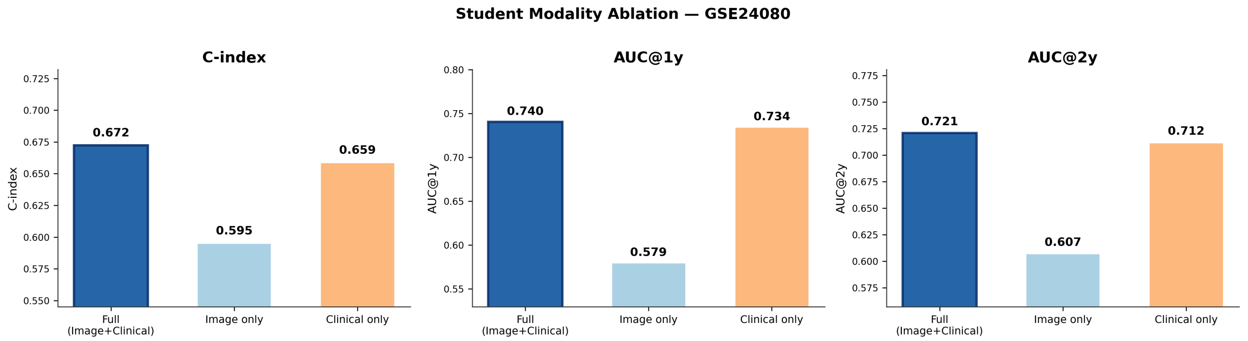


**Figure S4:** Modality ablation for the distilled student model on the external GSE24080 cohort ($t = 1$ month, $n = 507$). C-index, $\text{tdAUC}_{1\text{yr}}$, and $\text{tdAUC}_{2\text{yr}}$ are shown for the full student model (DeepInsight image + five clinical features), image-only, and clinical-only configurations. Full: C-index = 0.672; Image only: 0.595; Clinical only: 0.659. Numerical values are provided in Supplementary Table S6.

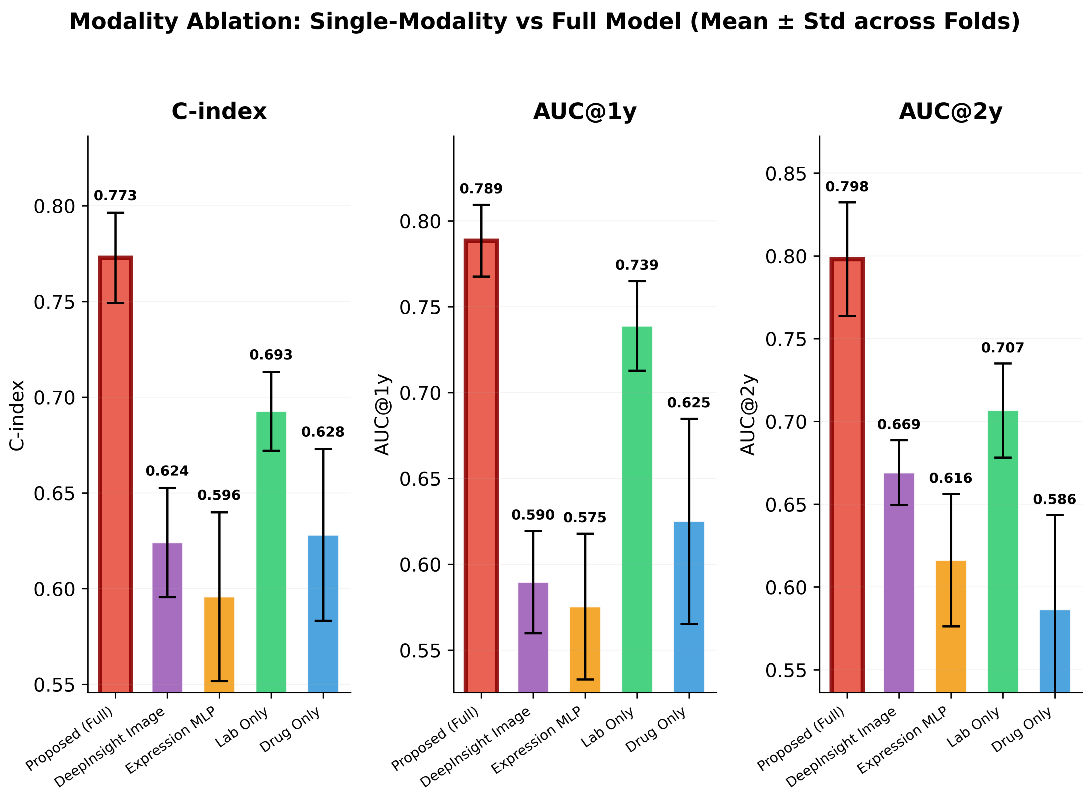


**Figure S5:** Internal modality ablation on the CoMMpass development set (five-fold CV, mean ± SD). C-index, $tdAUC_{1yr}$, and $tdAUC_{2yr}$ are shown for each single-modality configuration and the full multimodal model. Error bars represent one standard deviation across five folds. Complements main-text Table 2.

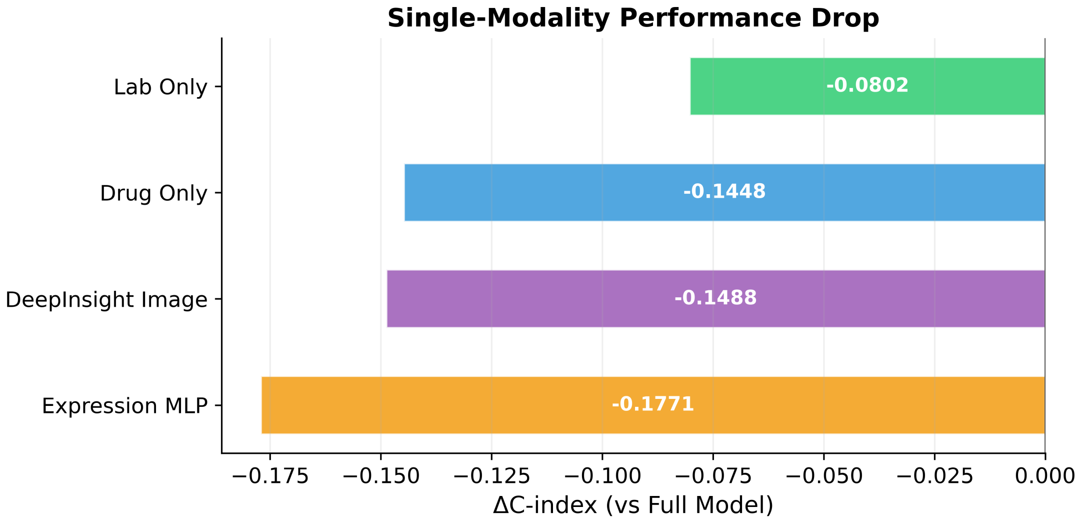


**Figure S6:** Single-modality performance drop relative to the full multimodal model, expressed as ΔC-index (five-fold CV mean). Each bar shows the reduction in C-index when the model is restricted to a single data source: laboratory trajectories only ($\Delta = -0.080$), drug history only ($-0.145$), DeepInsight image representation only ($-0.149$), and expression MLP baseline ($-0.177$). Longitudinal laboratories exhibit the smallest performance drop, consistent with their central role in ISS/R-ISS staging.

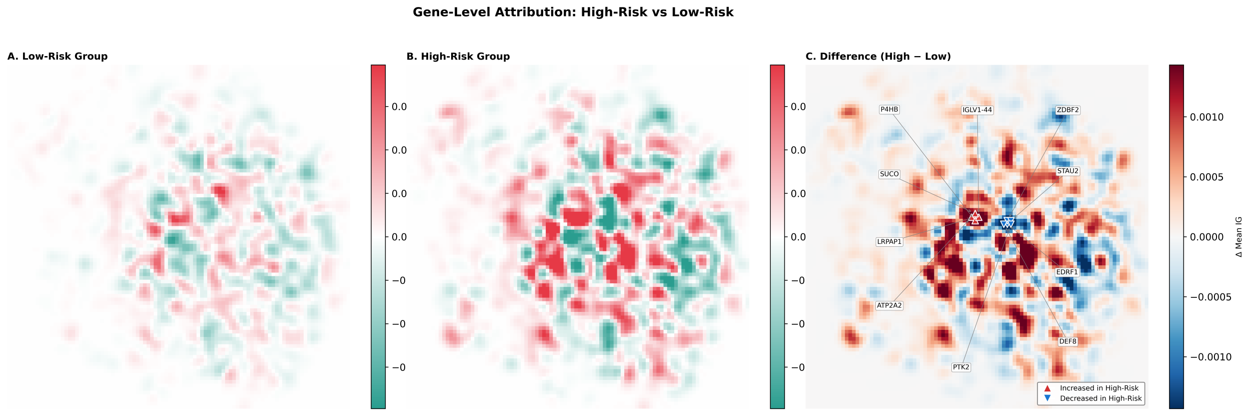


**Figure S7:** Representative Integrated Gradients attribution maps on the DeepInsight gene expression images, grouped by model-predicted risk category. **(A)** Low-risk patients exhibit diffuse, low-magnitude attribution patterns. **(B)** High-risk patients show spatially concentrated, high-magnitude activations. **(C)** Difference map (high-risk minus low-risk) highlighting genomic regions with the largest risk-associated attribution shift; individual genes are annotated at peak locations by reverse-mapping pixel coordinates to gene identifiers via the pyDeepInsight coordinate matrix.

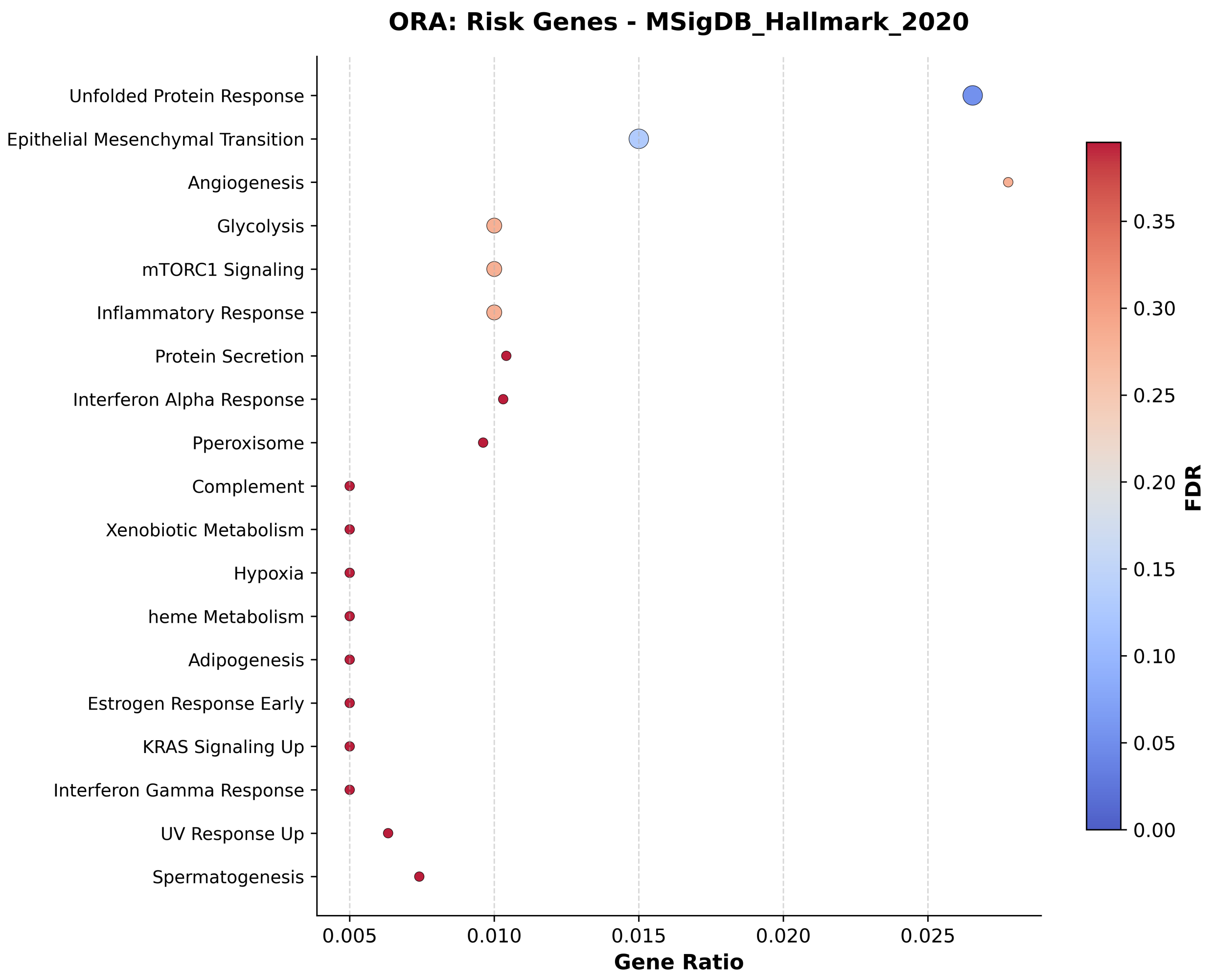


**Figure S8:** Over-representation analysis (ORA) of risk-increasing genes against the MSigDB Hallmark 2020 gene set collection. Risk-increasing genes were defined as those with positive mean Integrated Gradients attribution (i.e., higher expression associated with higher predicted log-hazard). Dot size is proportional to the gene ratio (number of overlapping genes / gene set size); colour indicates false discovery rate (FDR). Top enriched terms include Unfolded Protein Response, Epithelial–Mesenchymal Transition, mTORC1 Signalling, Glycolysis, and Angiogenesis.

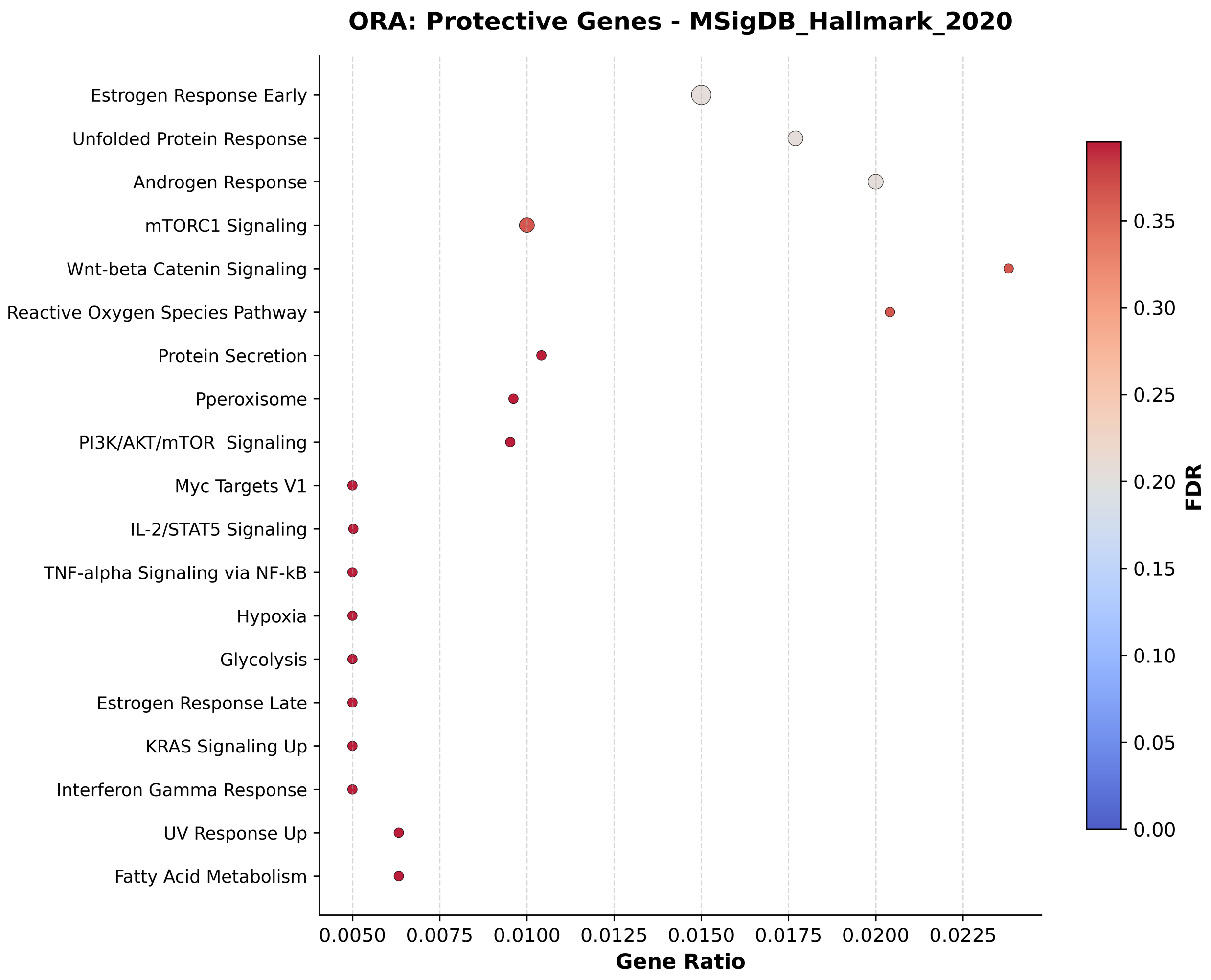


**Figure S9:** Over-representation analysis (ORA) of protective genes against the MSigDB Hallmark 2020 gene set collection. Protective genes were defined as those with negative mean Integrated Gradients attribution (i.e., higher expression associated with lower predicted log-hazard). Top enriched terms include Estrogen Response Early, Unfolded Protein Response (with gene members distinct from those in Figure S8), Androgen Response, and Wnt/$\beta$-Catenin Signalling. The presence of UPR in both risk and protective sets reflects the dual role of endoplasmic reticulum stress in plasma cell biology.

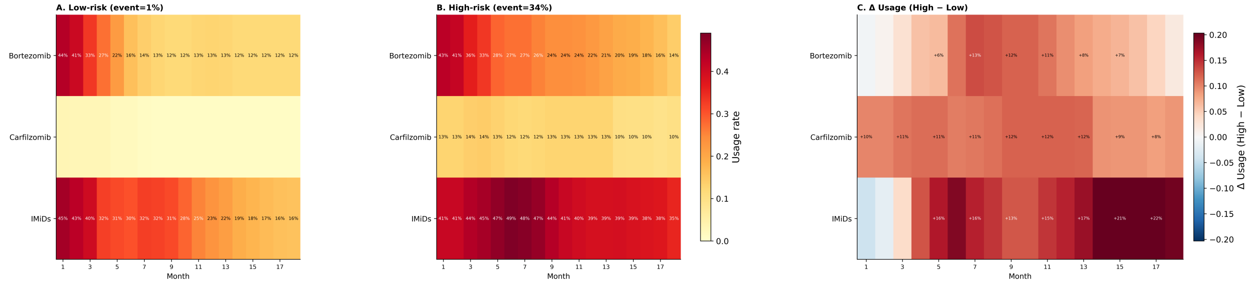


**Figure S10:** Drug utilisation patterns stratified by model-predicted risk group on the held-out CoMMpass validation set. Monthly administration rates are shown for bortezomib, carfilzomib, and immunomodulatory drugs (IMiDs). The high-risk group (death rate 34%) received higher carfilzomib and IMiD utilisation across all time bins compared with the low-risk group (death rate 1%), consistent with intensified treatment for higher-risk disease. These patterns reflect indication bias inherent in observational treatment data rather than causal treatment effects.

### Supplementary Methods

#### Evaluation protocol note

The 624-patient CoMMpass development set was partitioned into five stratified folds for model training. Within each fold, checkpoint selection and early stopping were performed using fold-internal validation metrics only. After training, the resulting fold-specific models were each evaluated on a separate common held-out CoMMpass validation split ($n = 128$), which was not used for parameter fitting within any fold. Cross-fold mean ± SD on this common held-out validation split are reported as the primary comparative performance summary (main-text Table 1; Supplementary Figure S2).

For downstream visualisation and representative held-out analyses, we additionally report a selected fold (fold 2), chosen by the highest mean $\text{tdAUC}_{1\text{yr}}$ across the primary landmarks ($t = 6, 12$ months) on the common held-out CoMMpass validation split. This selected fold was used for the landmark-wise plots, Kaplan–Meier stratification figures, and teacher-to-student transfer analyses (main-text Figures 2–5; Supplementary Tables S2–S6).

For benchmark transfer to the external GSE24080 cohort, each benchmarked method was represented by its own fold-specific model selected on the same common held-out CoMMpass validation split. The external cohort was used strictly for final external evaluation only; no model selection, fold selection, checkpoint selection, threshold derivation, or hyperparameter tuning used any GSE24080 data.

#### Hyperparameter configuration

**Optimisation.** Training used AdamW optimisation with a base learning rate of $1 \times 10^{-4}$, weight decay 0.01, and linear warmup over 30 epochs. A ReduceLROnPlateau scheduler (factor 0.5, patience 15 epochs) reduced the learning rate upon metric stagnation. Gradient norms were clipped at 1.0. Batch size was 48 with gradient accumulation over 4 steps, yielding an effective batch size of 192; this configuration was chosen to ensure that each Cox partial likelihood batch contains a sufficient number of events for stable gradient estimation despite the low overall event rate (19.9%).

**Early stopping and checkpoint selection.** Early stopping monitored the EMA-smoothed ($\alpha = 0.35$) validation $\text{tdAUC}_{1\text{yr}}$ averaged across primary landmarks ($t = 6, 12$ months), with patience 80 epochs and a minimum epoch gate of 100 per fold (maximum 600 epochs per fold). The minimum epoch gate ensures that the curriculum-weighted $t$-sampling has completed its transition from early-landmark concentration to balanced coverage before checkpoint selection begins. Model weight EMA (Polyak averaging, decay 0.998) was used for validation and final evaluation.

**Sampling and curriculum.** Patient-level sampling drew $K = 3$ (patient, $t$) pairs per patient per epoch. For each sample, the landmark month $t$ was assigned by a curriculum-weighted distribution: during the first phase of training, the distribution concentrates probability on early landmarks where event density is highest, then transitions linearly toward uniform coverage across $t \in \{1, \ldots, 18\}$ as training progresses. This curriculum ensures that the model first learns from the most informative risk-set compositions before extending to sparser later landmarks.

**Augmentation.** Image augmentation consisted of Gaussian noise ($\sigma = 0.02$) and random erasing ($p = 0.1$). Geometric augmentations (flips, rotations) were disabled because the $t$-SNE-derived spatial layout of the DeepInsight image lacks natural rotational or reflective symmetry—flipping or rotating the image would destroy the co-expression neighbourhood structure that the CNN is designed to exploit.

**Regularisation.** Deploy-aware modality dropout (probability 0.30) jointly zeroed laboratories and drugs during training; the image modality was never dropped. Auxiliary head weights: image $\lambda_{\text{img}} = 0.15$, tabular $\lambda_{\text{tab}} = 0.10$.

**Architecture details.** Lightweight CNN: 5 convolutional blocks ($3 \times 3$ conv, BN, ReLU, $2 \times 2$ max-pool), adaptive average pooling, FC projection (dropout 0.3). Parameters: $\sim$1.0M. Image encoder learning rate multiplier: 0.1. Dual-stream lab Transformer: 2 layers, 4 heads, $d = 128$, FFN = 256. Drug Transformer: 2 layers, 4 heads, $d = 64$. Gated fusion: gate hidden 64, gate dropout 0.30, fusion dropout 0.35. Time embedding: $d = 16$. Fusion hidden: $d = 128$.

#### Risk stratification threshold derivation

For Kaplan–Meier risk stratification, predicted risk scores at each landmark were dichotomised into high- and low-risk groups. The threshold at each landmark was determined by maximising the log-rank test statistic over a grid of candidate cut-points applied to the training-set risk score distribution. These training-derived thresholds were then applied without modification to the held-out validation set and the external cohort (GSE24080), ensuring that no information from the evaluation data influenced the group assignment.

#### Knowledge distillation configuration

Student input: DeepInsight image ($96\times96\times1$, mutation channel zeroed) + 5 clinical features (HGB, CREAT, ALB, LDH, $\beta_2$M). Architecture: LightweightCNN + clinical MLP; $\sim$1.0M parameters. Image encoder initialised from teacher. Loss weights: $\alpha_{\text{surv}} = 0.3$, $\alpha_{\text{out}} = 0.5$, $\alpha_{\text{feat}} = 0.2$. Learning rate: $3 \times 10^{-4}$. Single landmark $t = 1$ month. Best checkpoint: epoch 86/146 (EMA tdAUC$_{\text{1yr}}$ = 0.831).
